# Genomic insights into bacterial kidney disease resistance in Arctic charr (*Salvelinus alpinus*) via a 72k SNP array

**DOI:** 10.64898/2026.06.25.734482

**Authors:** Christos Palaiokostas, Henrik Jeuthe, Katja Nilsson, Hampus Hällbom, Charlotte Axén, Øystein Evensen, Susanne Eriksson, Martin Johnsson

## Abstract

Selection for disease resistance forms one of the most highlighted areas of aquaculture breeding. A breeding program for Arctic charr has been operating in Sweden for over 40 years, making it the oldest of its kind worldwide for this species. However, the lack of available genomic resources prevented selection for any disease-resistance traits. A 72k Axiom SNP array was produced in this study and used to assess the potential to select for charr resistant to bacterial kidney disease (BKD), which is currently a major threat to the industry. Following a challenge experiment with *Renibacterium salmoninarum*, the causative agent of BKD, relevant phenotypic proxies were collected from approximately 2,000 charr. Thereafter, those animals were genotyped with the new 72k SNP array. The magnitude of the estimated variance components suggested potential for breeding for BKD resistance in charr, with relevant heritabilities ranging from 0.05 to 0.56 depending on the resistance proxy used. In addition, GWAS suggested that BKD resistance is a polygenic trait. Furthermore, genomic prediction approaches indicated that BKD-resistant animals can be identified using their SNP genotypes. Accuracies, expressed as Pearson correlation coefficients, when BKD resistance was analysed as a continuous trait, ranged from 0.42 to 0.52. In the scenario where BKD resistance was treated as a binary trait, the efficiency of genomic prediction was assessed using ROC curves, with an area under the curve of 0.72. Finally, no unfavourable correlations were found with growth traits. The developed 72k SNP array has the potential of being a pivotal tool for the Swedish Arctic charr breeding program. Moreover, our data support the use of genomic prediction in breeding BKD-resistant Arctic charr. As a critical next step, further validations in actual industry conditions would be required.

## Introduction

Bacterial kidney disease (BKD), caused by *Renibacterium salmoninarum*, poses a significant threat to salmonid farming (Bayliss et al. 2018), amongst which Arctic charr (*Salvelinus alpinus*) is particularly vulnerable. In Sweden, BKD is considered the most damaging disease for the domestic charr industry, with recorded mortalities reaching up to 80% (Andersson et al. 2023). The lack of effective therapeutic agents exacerbates the problem, with the most effective way to control the infection being to slaughter the animals and disinfect the facility (Persson et al. 2022). Breeding for genetically resistant animals might offer solutions and mitigate the problem (Huang et al. 2025).

Considerable effort has been devoted to selecting for disease resistance across various aquaculture species (Robinson et al. 2023). This is hardly surprising, as the most common production system in aquaculture, which uses net pens in lakes or the sea, entails constant interaction with the surrounding aquatic environment. As such, any treatment directly affects the ecosystem, limiting the pharmaceuticals available. Furthermore, as aquaculture is a relatively recent industry, effective therapeutic agents remain scarce for many diseases (Kowalska et al. 2020).

The adoption of genomics in aquaculture breeding has propelled progress, enabling the detection of genes associated with genetic resistance (Pavelin et al. 2021; D’Ambrosio et al. 2025; Mukiibi et al. 2025; Barría et al. 2026) and the implementation of genomic selection (Meuwissen et al. 2001) in commercial breeding programs (Vallejo et al. 2019; Griot et al. 2021; Yáñez et al. 2023). As is typical of traits related to disease resistance, the breeding candidate *per se* does not have a phenotype, and selection relies on information from relatives. The inclusion of genomic information in such cases has repeatedly been shown to substantially increase selection accuracy compared to traditional pedigree-based approaches (Villanueva et al. 2011; Palaiokostas et al. 2016; Oikonomou et al. 2022; Vela-Avitúa et al. 2022; Vallejo et al. 2024; Guddanti et al. 2026; Liang et al. 2026).

Similar to livestock, single-nucleotide polymorphism (SNP) arrays are the tool of choice for modern breeding practices in aquaculture species. Even though genotype-by-sequencing approaches have been shown to be valuable (Robledo et al. 2018), the “cleaner” genotypes obtained from SNP arrays, together with the more straightforward downstream analysis requirements, largely explain their widespread adoption in breeding.

A national breeding program for Arctic charr has been operating in Sweden for over 40 years, focusing primarily on growth-related traits (Palaiokostas et al. 2021). Notably, no trait related to disease resistance has previously been included in the program’s breeding goal. To the best of our knowledge, the potential to select for BKD resistant charr is unknown, as no prior heritability estimates are available. Notably, a moderate heritability of approximately 0.23 has been reported for Atlantic salmon (*Salmo salar*) (Gjedrem and Gjøen 1995).

Until recently, the limited availability of genomic resources has restricted the implementation of genomic-based breeding decisions. As such, the Arctic charr program has been relying entirely on pedigree-derived breeding values. The recent development of a high-density Axiom array (Thermo Fisher Scientific^™^) containing approximately 600,000 SNPs (Palaiokostas and Johnsson 2026) has been pivotal for the future of the breeding program, as it enables routine use of genomic selection.

From a global perspective, Arctic charr is more of a "niche" aquaculture species, and even though it is the second most common fish species in Swedish aquaculture, its annual production does not exceed 1,500 tonnes. Thus, the routine use of a high-density array is financially challenging. Previous studies in various aquaculture species have shown that genomic selection can be successfully implemented with relatively low SNP densities (Palaiokostas et al. 2019; Kriaridou et al. 2020; Fraslin et al. 2023; Kriaridou et al. 2023). Since SNP density is one of the factors that determine genotyping costs, lower densities would reduce per-sample costs without compromising selection efficiency (Liu et al. 2024). Naturally, the above also depends on the actual underlying genetic architecture (Song et al. 2023). Regardless, linkage disequilibrium profiles previously obtained from genotyping charr from various year classes of the breeding programs using the high-density array suggested that a low- to medium-density array would be suitable for genomic selection (Palaiokostas and Johnsson 2026).

In the current study, a medium-density array was developed containing 72k SNPs selected from the aforementioned high-density one. Approximately 2,000 juvenile Arctic charr from the 2025 year-class of the Swedish breeding program were phenotyped for proxy indicators related to BKD resistance. Those phenotypes were derived from a disease challenge experiment. Thereafter, those animals were genotyped with the 72k SNP array. Heritability estimates for the resistance proxies were estimated, followed by genome-wide association analysis (GWAS). Finally, we investigated the potential of using genomic selection to identify genetically resistant animals.

## Methods

### Medium-density array development

The selection of SNPs for the high-density Arctic charr array has been previously described (Palaiokostas and Johnsson 2026). Subsequent filtering of those markers for developing a medium-density array involved a minimum call rate of 99%, a minimum minor allele frequency of 0.15, and no statistically significant deviations from Hardy-Weinberg equilibrium (α = 0.001). Moreover, only SNPs with a minimum physical distance of 5 kbp between them were selected. Following further recommendations from Thermo Fisher Scientific, a subset including 72k of the above SNPs was tiled in an Axiom array, which was trademarked as *Salpinus*. Those recommendations were derived from their proprietary software, which provides a probabilistic estimate of the likelihood that a given SNP will be converted into a reliable SNP assay.

### Background of the studied population

Our study used Arctic charr from the Swedish national breeding programme (Carlberg et al. 2018). The base population originated from wild fish collected in 1979 in Lake Hornavan, northern Sweden. The initial number of founding animals is unknown, with recent estimates suggesting an effective population size (N_e_) of approximately 20 (Palaiokostas and Johnsson 2026). The programme operates within a closed breeding nucleus, with discrete generations, using a 1:2 sire-to-dam mating ratio. The number of families in each generation has ranged from 45 to 125. Currently, the 10th generation of the breeding programme has been formed, comprising 89 full-sib families.

### BKD challenge experiment and phenotypic recordings

The bacterial strain used for the disease challenge experiment was the SVA9 (4/86). It originated from one of the early Swedish BKD outbreaks and has been utilised in previous studies (Jansson et al. 1996). The bacterium was initially cultivated on KDMC agar plates. Colonies were harvested and transferred to KDM broth, then incubated on a shaking platform at 15°C. The OD was monitored with a PV4 visible spectrophotometer until the broth reached an estimated bacterial concentration of 5×10^7^ CFU/ml.

Juveniles from the 10th generation of the Swedish Arctic charr breeding program were transferred to VESO Aqualab in Norway during June 2025. In total, 2,200 fish (2-5 g) from 89 full-sib families were used. The animals were acclimated over a growth period of approximately two months, during which they reached an average weight of ∼20g. Thereafter, 2000 of those animals were infected through intraperitoneal (IP) injection with the prepared bacterium inoculum (0.1 ml per animal). The injected animals were split equally between two tanks (each containing 1000 animals) and monitored daily in accordance with VESO Aqualab’s standard operating procedures. Those involved continuous monitoring of water quality parameters, collection of dead fish, and observation of abnormal behaviour in the challenged animals (e.g., reduced swimming capacity, loss of appetite). The freshwater temperature was 12 °C (±1 °C), with a water flow of 0.8 L/kg/min, while a photoperiod of 24:0 (L:D) was used throughout the study.

During the experiment, fin-clips were collected from all fish for genotyping. Furthermore, head kidney tissue was collected from 800 animals for RT-qPCR analysis to investigate bacterial load in the fish. Those animals included approximately 400 that were among the first mortalities, while the rest survived until near the end of the experiment. The RT-qPCR analysis was performed at PatoGen’s facilities in Ålesund, Norway, using their accredited protocols (Norwegian Standard NS-EN ISO/IEC 17025). Ct values, representing the cycle number where fluorescence surpasses the detection threshold, were produced for each sample as indicators of a bacterial load. In general, the higher the initial bacterial load in a sample, the fewer PCR cycles are needed to produce a detectable amount of PCR product.

Overall, the following BKD resistance proxies were used for downstream analysis: days of survival (continuous trait), overall survival, and N_50_ survival (binary traits). Finally, for the subset of animals from RT-qPCR, the Ct values (a continuous trait) were analysed as a proxy for BKD resistance.

### Genotyping and quality control

Genotyping was performed in all 2000 charr juveniles that were challenged with *Renibacterium salmoninarum*. Genomic DNA was extracted from collected fin-clips, followed by genotyping with the new custom Axiom™ SNP array. Both activities were performed by Identigen (Dublin, Ireland). The obtained genotypic data were filtered using preGSf90 v1.26 from the BLUPf90 suite (Misztal et al. 2018), discarding SNPs with a call rate below 98% and minor allele frequency (MAF) below 0.05. Moreover, the same software was used to filter SNPs for which the number of observed heterozygotes deviated from the number expected under the Hardy-Weinberg equilibrium by more than 0.15 (Wiggans et al. 2009). Finally, the existence of any underlying population structure was assessed using principal component analysis (PCA) on the genomic relationship matrix (GRM). The GRM was estimated with preGSf90 v1.26 from the BLUPf90 software suite, following VanRaden (2008).

### Parentage assignment and pedigree reconstruction

During transport, all the animals were pooled in the same tank container. Because the fish were not tagged, each individual’s pedigree was therefore unknown. A subset of candidate parents had been previously genotyped (n = 96) with the high-density array (Palaiokostas and Johnsson 2026). Those genotypes were subsetted for the SNPs of the current array. Parentage assignment was performed using the seekparentf90 v1.59 from the BLUPF90 suite (Misztal et al. 2018), allowing for a genotypic error of 2%. For verification purposes, parentage assignment was also performed using the R package APIS v2.0.8 (Griot et al. 2020; Roche et al. 2024). In parallel, a “naive” approach to reconstructing the pedigree was performed using the GRM estimated with preGSf90 v1.26 from the BLUPf90 software suite, following VanRaden (2008). More specifically, only the off-diagonal elements of the GRM whose normalised values (each value was divided by the square root of the product of the corresponding diagonal elements; this was performed to obtain realised relationships) fell within the range 0.45 - 0.65 were retained as indicative of potential full-sibs. Thereafter, a network analysis was conducted using igraph v2.2.1 (Csárdi et al. 2026), and the resulting clusters indicated full-sib families (https://github.com/chpalaiokostas/Genetic_resistance_BKD/tree/main/Pedigree_reconstruction).

### Estimation of variance components

As phenotypic proxies of BKD resistance, we used the number of surviving days both as a continuous and a right-censored trait, overall survival, N_50_ survival and the Ct values from RT-qPCR. Variance components for those resistance proxies were estimated with blupf90+ v2.73 and gibbsf90+ v3.24 from the BLUPf90 suite (Misztal et al. 2018) using the following animal model:

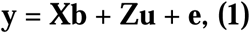

where **b** is the vector of the fixed effects (intercept, tank); **X** is the incidence matrix relating phenotypes with the fixed effects; **Z** is the incidence matrix relating phenotypes with the random animal effects; **u** is the vector of random animal effects ∼ N(0, **G**σ_g_^2^) where **G** is the GRM (VanRaden 2008), σ_g_^2^ is the additive genetic variance; **e** the vector of residuals ∼N(0, **I**) and σ_e_^2^ is the residual variance. The heritability was estimated using the following formula:

For the overall and N_50_ survival, the probit link function was used to map the observed binary phenotype (0 = Non-resistant, 1 = Resistant) onto the underlying liability scale. A residual variance on the underlying scale is not identifiable in threshold models (Goldstein et al. 2002) and was therefore fixed to 1.

The parameters of each of the above models were estimated through Markov chain Monte Carlo (MCMC) using Gibbs sampling. Each MCMC involved 1 M iterations, out of which the first 100,000 were used as burn-in. Finally, a thinning step of 1,000 was used.

### Genome-wide association analysis

A weighted genomic best linear unbiased prediction (WGBLUP) analysis was performed (Zhang et al. 2016; Lourenco et al. 2020) using the BLUPF90 software suite, as previously described (Palaiokostas et al. 2022). In short, a genomic relationship matrix (GRM) was estimated following VanRaden (2008) as:

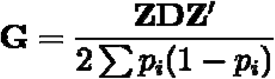

where **Z** *is* a matrix of centered genotypes, **D** is a weight matrix for all SNPs, and p_i_ is the corresponding MAF of each SNP. Thereafter, SNP weights were calculated using the nonlinear A method (VanRaden 2008). The above process was followed for two iterations as previously described (Palaiokostas et al. 2022). Finally, the percentage of explained additive geneti variance was estimated using non-overlapping windows of adjacent SNPs, each covering a physical distance of 1 Mbp.

### Genomic prediction for BKD resistance

An assessment of the efficiency of predicting BKD resistance in charr based on their SNP genotypes was conducted. Different approaches were used depending on whether the resistance proxy was the number of days of survival or a binary outcome. The dataset was split into training and validation/test sets. The size of the splits varied depending on the BKD resistance proxies used, as described below. Overall, SNP effects were estimated from direct genomic values (DGVs) derived from GBLUP (Lourenco et al. 2020). Thereafter, those SNP effects were used to estimate DGVs for animals in the test sets where the phenotypes had previously been masked.

For the surviving days, a three-fold cross-validation approach was used. More specifically, the dataset was split into sequential training (n=1337) and validation sets (n=667). In the validation sets, the animals’ phenotypes were masked, and their DGVs were estimated using the SNP effects derived from the training set. This cross-validation procedure was repeated 5 times to minimize potential bias arising from the random allocation of animals to the training and validation sets. The blupf90+ v2.73 and the postGSf90 v1.89 were used to estimate GEBVs and SNP effects, respectively, in the training sets. DGV predictions for the test sets were performed using predf90 v2.04 from the BLUPf90 suite (Misztal et al. 2018). The accuracy of the DGVs from the validation set was assessed using the Pearson correlation coefficient between them and the number of surviving days.

In the case where BKD resistance was studied as a binary trait (N_50_ survival), SNP effects were estimated from DGVs derived from animals of a single tank (n=1000). Therefore, half of the dataset (animals from either of the two tanks) was used for training, with the other half forming the test set. This approach was repeated for both tanks. Since the animals were randomly allocated to the two tanks during the challenge experiment, no biases due to unequal family representation were expected. The gibbsf90+ v3.24 and the postGSf90 v1.89 were used to estimate GEBVs and SNP effects, respectively, in the training sets. In the case of Gibbs sampling, the MCMC parameters involved 1 M iterations, of which the first 100,000 were used as burn-in. In addition, a 1,000-iteration thinning step was included. As in surviving days, the predf90 v2.04 from the BLUPf90 suite (Misztal et al. 2018) was used to estimate GEBVs for the animals in the test set. The assessment in this case was initially performed using a “naive” approach, in which animals from the test set with a DGV > 0 were classified as resistant. Equally, animals with a negative DGV were classified as non-resistant. A confusion matrix was computed to assess the model’s classifications against the actual phenotype. At the same time, receiver operator characteristic (ROC) curves were also used to assess the efficacy of classifying charr as resistant or non-resistant (based on N_50_ survival). The area under the curve (AUC) (Hanley and McNeil 1982; Wray et al. 2010) was used to assess performance, with a value of 1 indicating a perfect classifier.

### Associations between growth and BKD EBVs

A preliminary analysis was conducted to investigate associations between EBVs for growth and BKD. Breeding values for BKD resistance (surviving days) were estimated for the identified parents of the challenged charr. Simultaneously, growth-related EBVs were available for those animals, as they were routinely estimated for the Arctic charr breeding program. Those EBVs were estimated using the blupf90+ v2.73 (Misztal et al. 2018). BKD resistance EBVs from 36 broodfish were regressed against each animal’s corresponding growth EBV. In addition, a Pearson correlation coefficient was estimated between growth and BKD resistance EBVs.

## Results

### Descriptive analysis of BKD resistance traits

The challenge trial was concluded after a six-week observation period. Mortalities began 16 days post-challenge and peaked during the fourth week (Figure 1). Overall, only ∼2% of the challenged population survived with no evident differences between the two tanks.

**Figure 1.**
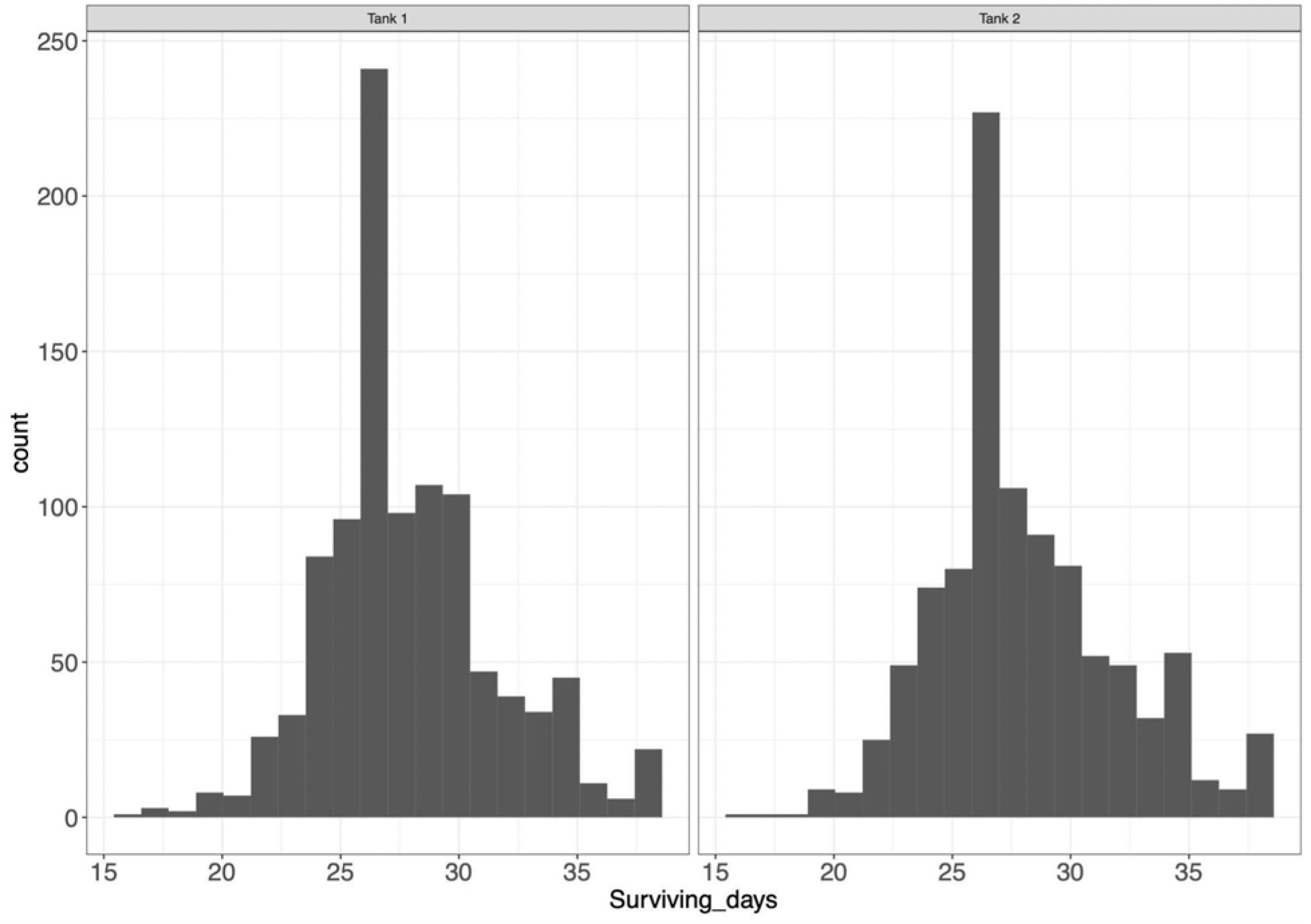
Surviving days of Arctic charr juveniles post-challenge intraperitoneal injection (IP) with an inoculum containing *Renibacterium salmoninarum* (5×10^6^CFU/fish). Following infection, the fish were randomly assigned to two tanks

A subset of the challenged animals, which either died early or survived towards the end of the experiment (n = 800), were sampled for RT-qPCR. No differences were observed in the obtained Ct values among the animals in the two tanks (Figure 2). The Ct values had a negative and statistically significant association with surviving days. A regression coefficient of −0.08 was estimated (SE 0.02; P < 10^-5^), while the Pearson correlation coefficient between Ct values and surviving days was −0.15 (P < 1-e04; 95% −0.22 to −0.08).

**Figure 2.**
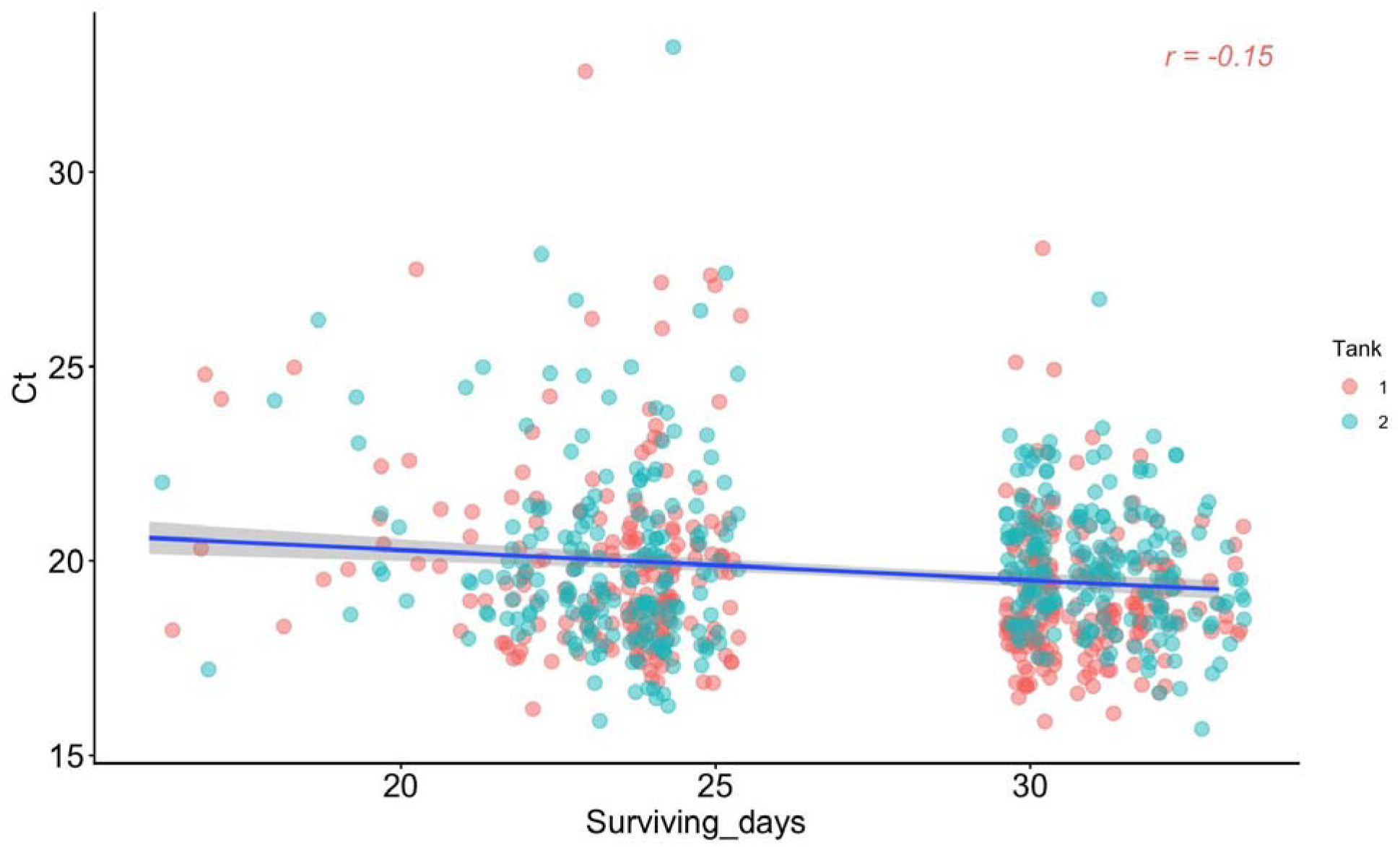
Scatterplot of Ct values obtained through RT-qPCR versus surviving days from Arctic charr juveniles challenged with *Renibacterium salmoninarum*. The plot depicts a regression line in blue, with 95% intervals in grey.

### Quality control of SNP genotypes

During QC, 18 animals were removed due to a call-rate below 90%. Moreover, 797 variants were removed due to MAF (< 0.05), 908 variants due to call-rate below 98%, and 31 variants were removed due to deviations from the Hardy-Weinberg equilibrium. Overall, 1982 animals and 69,745 SNPs were retained for downstream analysis. The PCA, as expected, did not reveal any obvious population structure with the first two principal components accounting for 2.8% and 2.5% of the observed variance, respectively (Figure 3).

**Figure 3.**
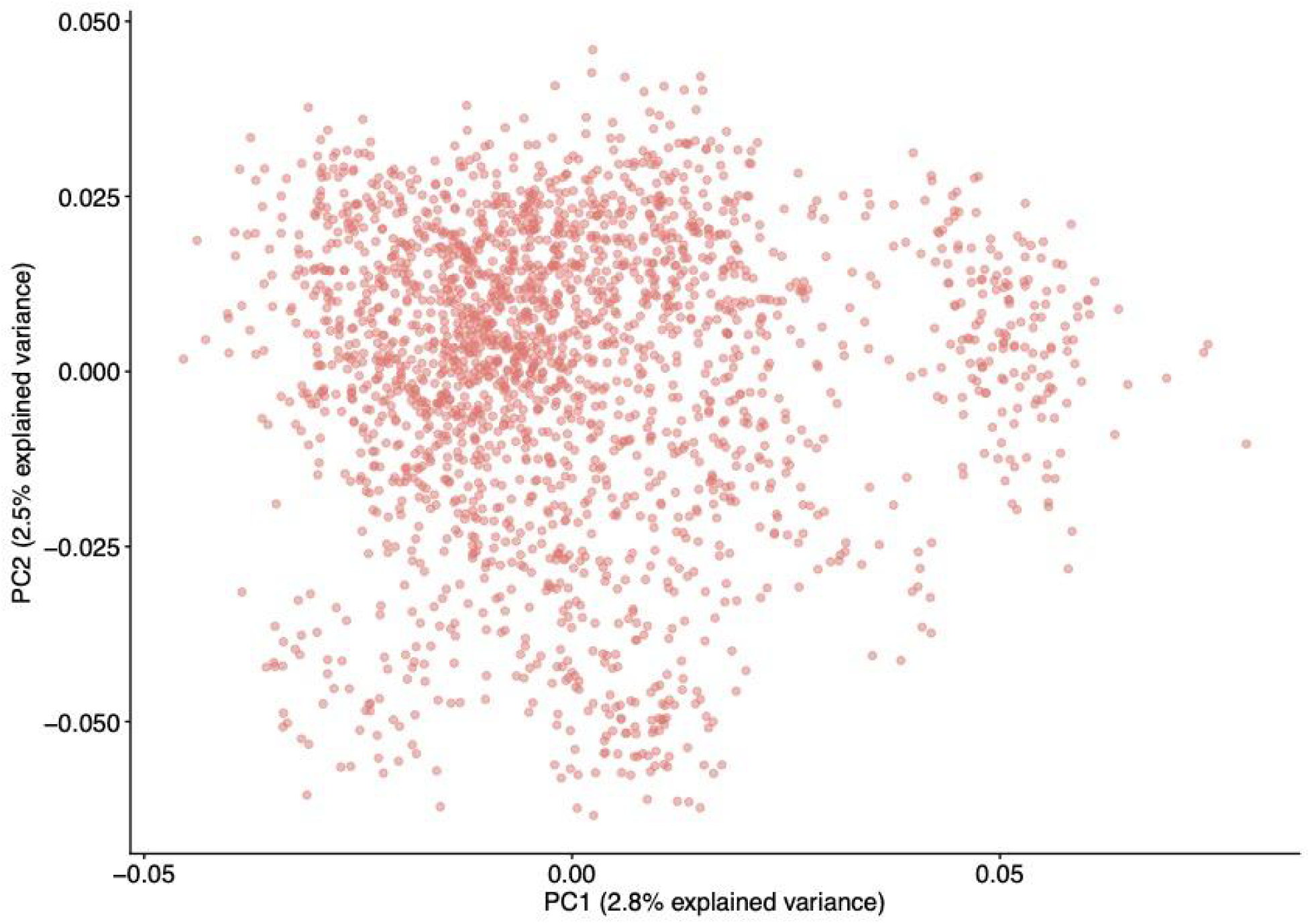
Principal component analysis of BKD-challenged Arctic charr juveniles (n = 1982). The PCA was performed on the genomic relationship matrix computed from 70,000 SNPs. The first two principal components accounted for 2.8% and 2.5% of the observed variation, respectively.

### Parentage assignment and pedigree reconstruction

Allowing for a maximum genotyping error of 2% in seekparentf90 resulted in 450 charr being assigned to unique parental pairs. These assignments were in full agreement with the ones from R/APIS. The animals uniquely assigned to a parental pair represented 18 full-sib families, each with 13 to 41 offspring. The mean size of those full-sib families was 24 offspring. In addition, 1,454 were assigned to at least a unique sire or dam. In the case of sires, 1032 charr were assigned, which represented 30 paternal half-sib families, each with 11 to 71 offspring. The mean offspring size of those half-sib families was 34. In the case of dams, 831 were assigned, representing 36 families, each with 13 to 41 offspring. Since each dam was mated with only one sire, those families were actually full-sib ones. The mean number of offspring in those families was 22. The network analysis conducted using the relationship values from the GRM (normalized values between 0.45 and 0.65) suggested the existence of 89 clusters (Figure 4), in agreement with the a priori expectation of the full-sib families used in the BKD challenge experiment.

**Figure 4.**
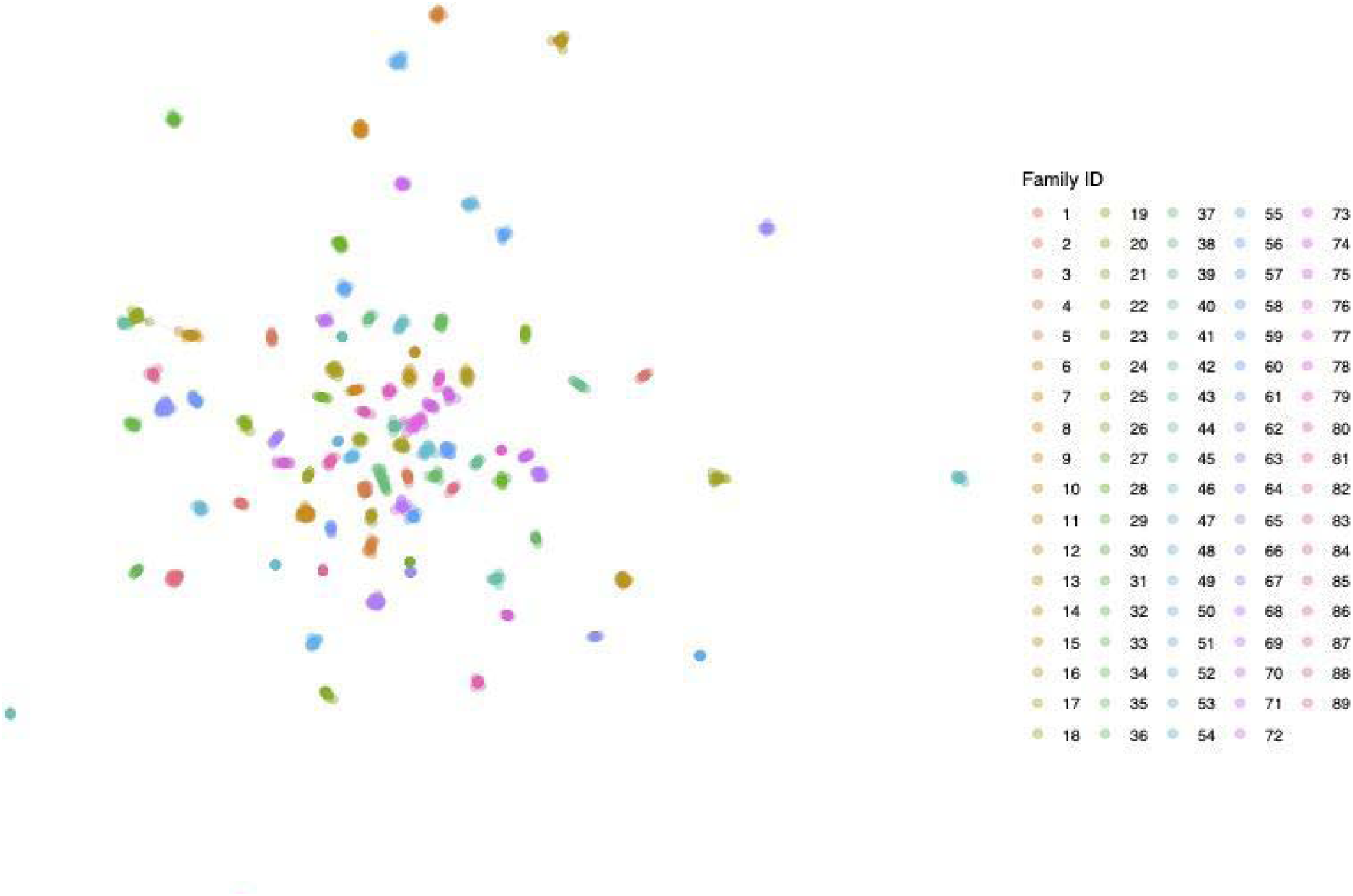
Pedigree reconstruction. Normalized genomic relationship values in the range of 0.45-0.65 were used for a network analysis conducted using the R package igraph v2.2.1. The suggested clusters indicate separate full-sib families.

### Estimation of variance components

The estimated heritabilities ranged from low to moderate, depending on the proxy used to characterise resistance to BKD (Table 1). In the case of Ct (related to bacterial load), where only a subset of animals were phenotyped (n=800), a low heritability estimate of 0.054 (SE 0.03) was found. Treating the number of surviving days as a proxy for resistance suggested a heritability of 0.46 (SE = 0.04). However, when the number of surviving days was treated as a right-censored trait the heritability dropped to 0.16 (SE = 0.03). On the other hand, when resistance was treated as a binary trait, heritabilities of 0.27 and 0.56 on the underlying scale were obtained for overall and N_50_ survival, respectively. Notably, especially in the former, the estimate was particularly imprecise, as the standard error exceeded half the estimate. Most likely, this was because one class (non-resistant) was overrepresented, as only 2% of the population survived the challenge experiment. In this case, the MCMC convergence appeared suboptimal, with an estimated autocorrelation coefficient of 0.5.

**Table 1.** Heritability estimates and accompanying standard error for proxies related to BKD resistance.

| Trait | Trait type | No. phenotypes | Software | Heritability | Standard Error |
| --- | --- | --- | --- | --- | --- |
| Surviving days | Continuous | 1982 | blupf90+ | 0.46 | 0.04 |
| Surviving days censored | Right censored | 1982 | gibbsf90+ | 0.16 | 0.03 |
| Overall survival | Binary | 1982 | gibbsf90+ | 0.27 | 0.15 |
| N<sub>50</sub> survival | Binary | 1982 | gibbsf90+ | 0.56 | 0.15 |
| Ct (RT-qPCR) | Continuous | 800 | blupf90+ | 0.054 | 0.038 |

### GWAS for surviving days and N_50_ survival

The GWAS results suggested that genetic resistance to BKD is a polygenic trait (Figure 5). Regions explaining just over 1% of the additive genetic variance were found in chromosomes 2, 8, 27, 28, and 35. A region on chromosome 2 accounted for the largest proportion of additive genetic variance, approximately 1.6%.

**Figure 5.**
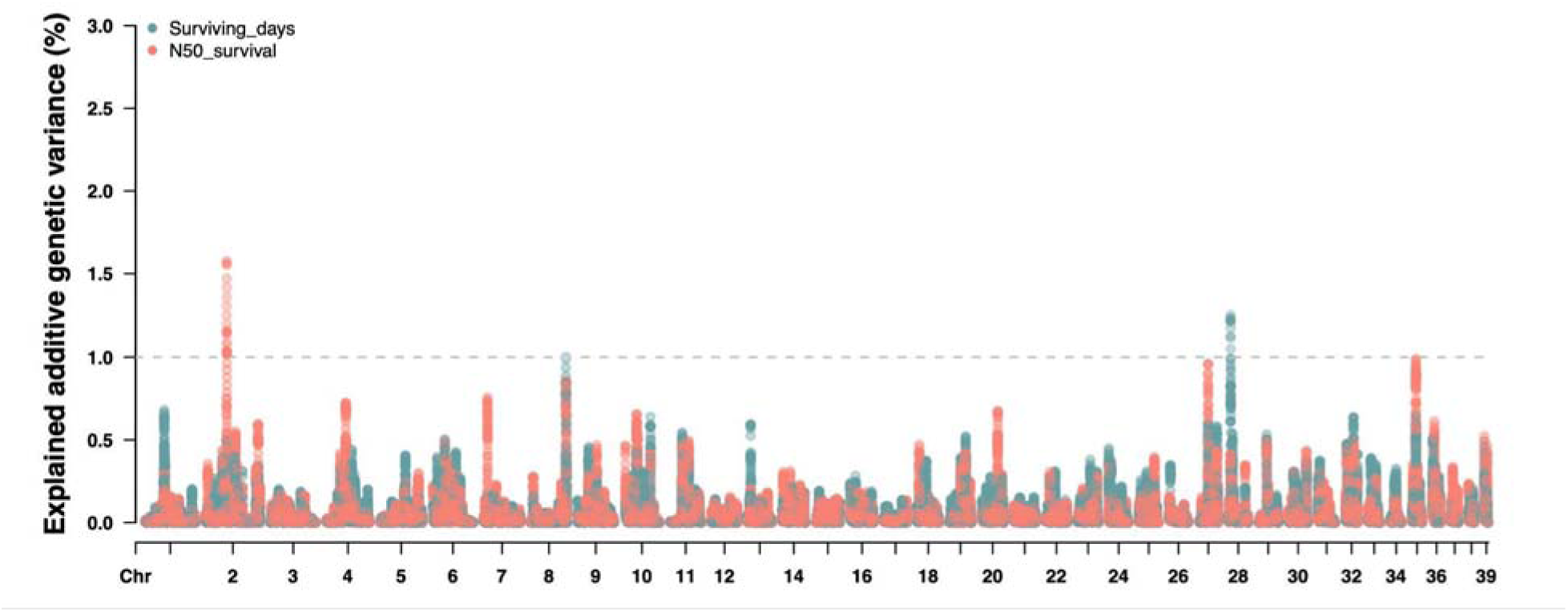
Manhattan plot showing the explained additive genetic variance for BKD resistance across the Arctic charr genome. The calculations were performed using WGBLUP with non-overlapping 1 Mbp windows. Two proxies of BKD resistance are depicted: Surviving days and N_50_ survival, each coded in a different colour.

### Prediction of BKD-resistant charr

At first glance, the use of genomic prediction appeared promising for selecting BKD-resistant charr. For surviving days, the accuracy, in terms of Pearson correlation, between the predicted DGVs of the validation sets and their corresponding phenotypes ranged from 0.41 to 0.52 (Table 2). The mean accuracy was 0.46, with an accompanying standard error of 0.01.

**Table 2.** Prediction accuracies of BKD surviving days using GBLUP.

| 3-fold CV | Replicate 1 | Replicate 2 | Replicate 3 | Replicate 4 | Replicate 5 |
| --- | --- | --- | --- | --- | --- |
| 1st fold | 0.48 | 0.47 | 0.45 | 0.45 | 0.48 |
| 2nd fold | 0.50 | 0.42 | 0.46 | 0.52 | 0.47 |
| 3rd fold | 0.43 | 0.44 | 0.45 | 0.49 | 0.45 |

The accuracies represent Pearson correlation coefficients between DGV and the number of surviving days. A 3-fold CV (cross-validation) strategy was performed, replicated 5 times.

For N_50_ survival, prediction efficiency was assessed by first inspecting boxplots showing the range of DGVs among resistant and non-resistant charr in the test set (each of the two tanks used in the challenge experiment was, in turn, used as the test set) (Figure 6A). In addition, the classification outputs were distinguished into true positives, true negatives, false positives, and false negatives (Figure 6B). Thereafter, classification was assessed more formally using a confusion matrix (Figure 6C). The correct classifications (the sum of true positives and true negatives) accounted for 64.4%. Regarding classification errors, false negatives accounted for 12.7% of predictions, while false positives accounted for 22.9%. As an overall metric for assessing classification performance, we used the AUC from the ROC curve. The AUC was 0.72. Finally, no obvious differences were found in any of the aforementioned metrics when either tank was used as a test set.

**Figure 6.**
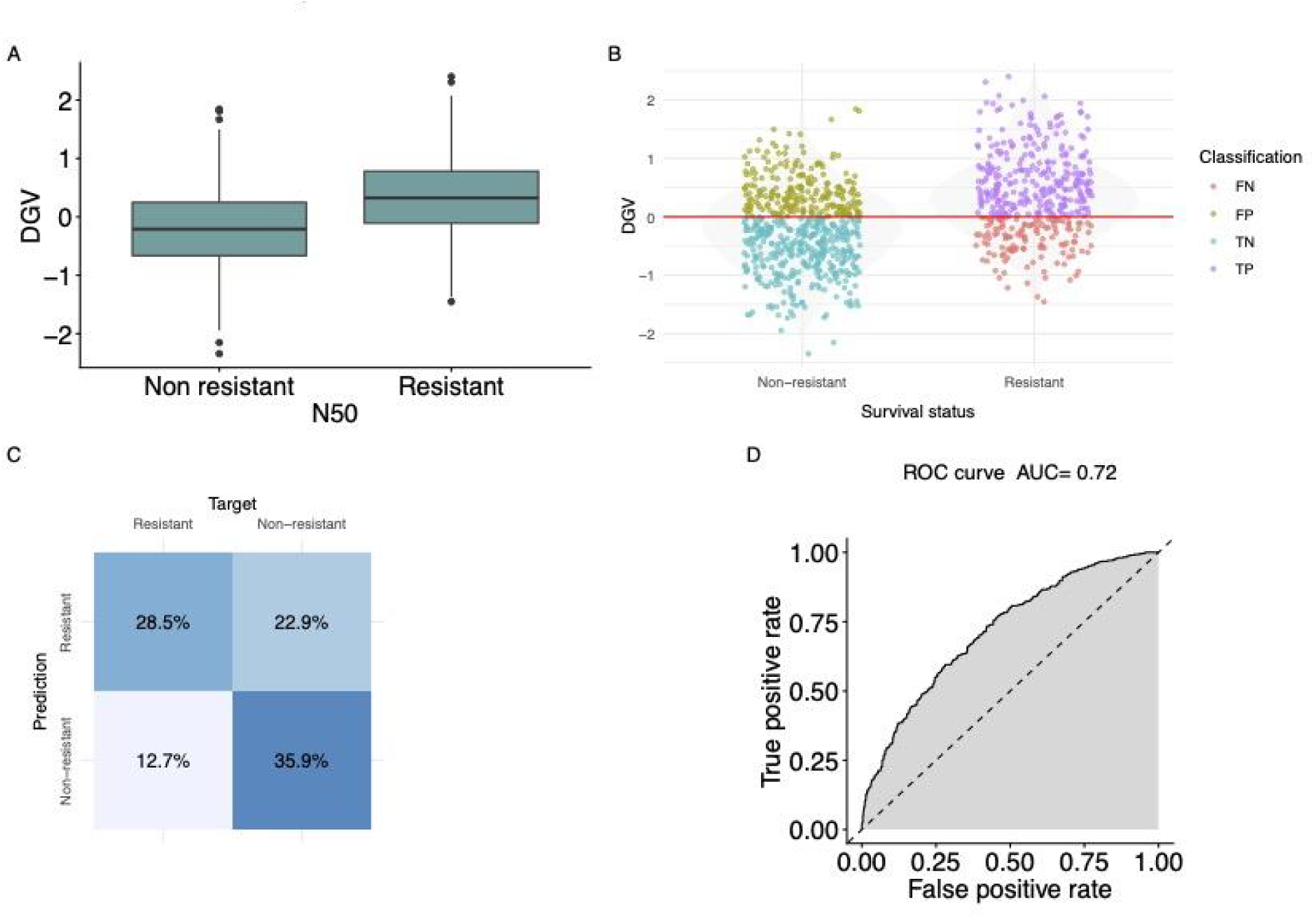
BKD resistance prediction analysis based on N_50_ survival. (A) Boxplot comparing the distribution of direct genomic values (DGV) between non-resistant and resistant groups. (B) A scatterplot depicting the classification categories in different colours versus the masked phenotype. (C) Confusion matrix displaying the classification performance of the prediction model, showing percentages for True Positives (TP; 28.5%), False Positives (FP; 22.9%), False Negatives (FN; 12.7%), and True Negatives (TN; 35.9%). (D) ROC curve and corresponding AUC metric

### Associations between growth and BKD EBVs

Only a weak, non-significant positive trend was observed between BKD resistance (measured in terms of surviving days) and growth EBVs. Specifically, regressing BKD resistance on growth EBVs yielded a regression coefficient of 0.012 (SE = 0.015; P = 0.43), while the Pearson correlation coefficient was 0.13 (Figure 7).

**Figure 7.**
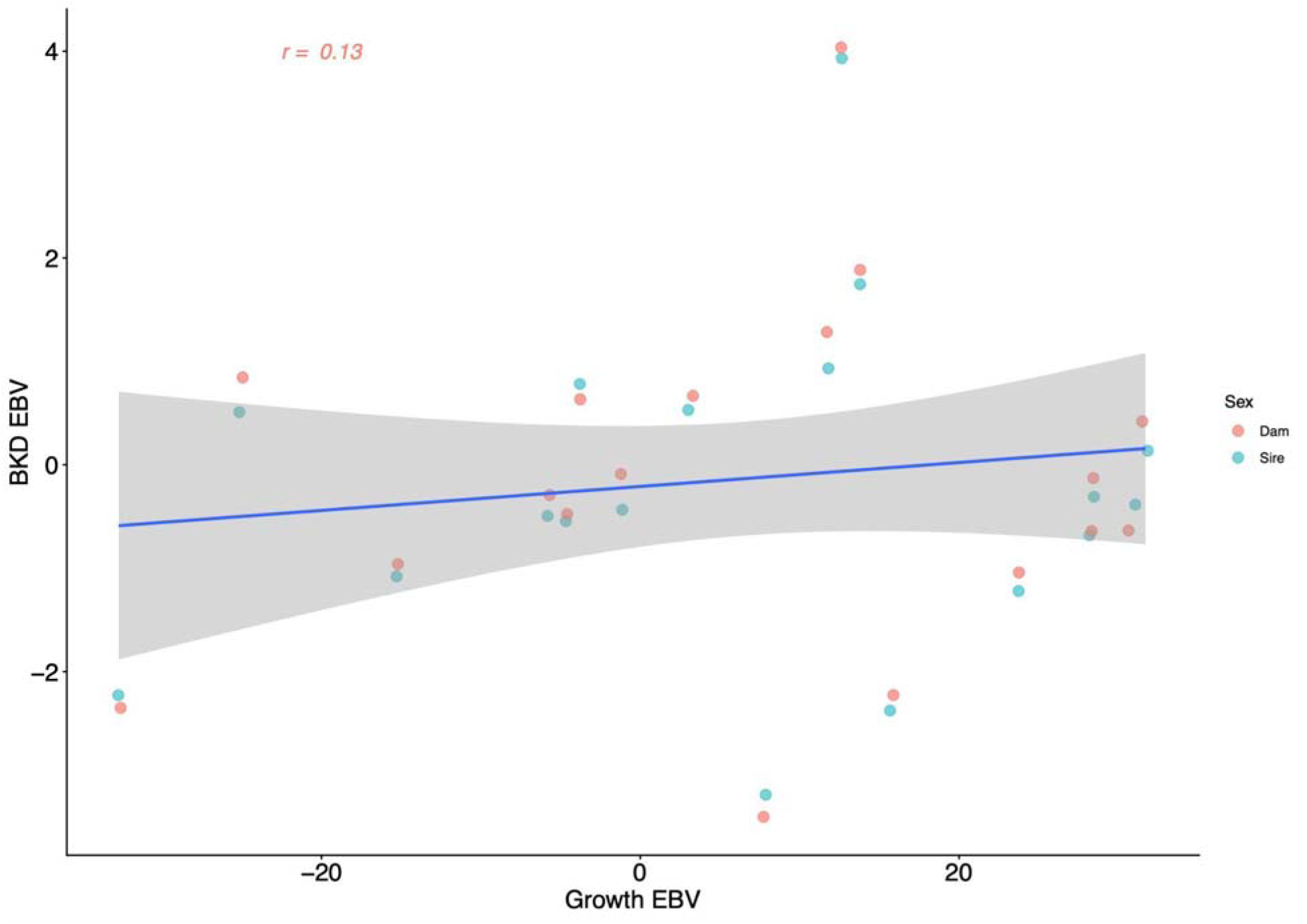
Relationship between BKD resistance and growth estimated breeding values (EBVs). The blue line represents the linear regression fit, with the grey shaded region indicating the 95% confidence interval. Individual data points represent the EBVs of single animals, colour-coded by sex (sires vs. dams).

## Discussion

A medium-density SNP array (72k) for Arctic charr was developed and used to assess the potential to select for BKD-resistant animals. As previously mentioned, the new array essentially contains a subset of the most informative and reproducible SNPs from a recently developed high-density array (600,000 SNPs) in the same population (Palaiokostas and Johnsson 2026). Genomic selection practices in animal breeding typically rely on medium-density SNP arrays (∼ 57k) as a more economically viable option. Densities of this magnitude also align with the relatively low N_e_ that characterizes many aquaculture breeding programs (Saura et al. 2021).

Our newly developed array appears well-suited to the Swedish Arctic charr breeding program. Amongst others, it circumvents one of the most highlighted weaknesses of SNP arrays, namely ascertainment bias when an array is developed in one population and used in others (Lachance and Tishkoff 2013). Critically, only a minimal number of SNPs were discarded (∼2.8%), even though strict thresholds were implemented using quality control (e.g., minimum call rate of 98%). For the size of the Arctic charr breeding program, a large number of animals, encompassing practically the entire number of families in the latest year class, were genotyped (∼2000). This high number of retained SNPs suggests that this array could play a pivotal role in the coming years by providing high-quality genomic data. In comparison, it is typical in salmonid breeding studies to discard 10% or more of the array’s SNPs, even with less stringent filtering thresholds (D’Ambrosio et al. 2019; Vallejo et al. 2019) or even when applied to the same population from which the array was initially produced (Dagnachew et al. 2025). Of course, to be more objective, it would be fair to claim that the newly developed array has a relatively narrow scope of applicability. Nevertheless, because information from other Arctic charr populations was included during array development, it is likely that our array would work well in other Nordic populations.

Our newly developed array was used here to assess the potential to select for BKD-resistant charr. An initial interpretation of the challenge trial results suggests that Arctic charr are particularly vulnerable to *Renibacterium salmoninarum*, with only approximately 2% of the challenged population surviving the six-week observation period. This high mortality rate appears to be in line with previously documented acute outbreaks in the species, where mortality reached up to 80% (Andersson et al. 2023). Since a controlled experimental setup was used here and the fish were injected with a bacterial inoculum, this discrepancy does not seem far-fetched. In contrast, previous studies in Atlantic salmon that also used an IP infection method reported lower mortalities ranging from 56% to 65% (Gjedrem and Gjøen 1995; Holborn et al. 2018). Those results suggest that Arctic charr might be more susceptible to BKD than Atlantic salmon. This also agrees with the fact that the bacterial cells injected here (5×10^6^ CFU/fish) were orders of magnitude lower than those previously used in Atlantic salmon (10^8^ CFU/fish) (Gjedrem and Gjøen 1995). However, as the fish of the aforementioned studies were substantially larger (40g and over 500g), such generalization might not be fully accurate. At the same time, it is possible that the bacterial strain in our study was more virulent than those previously used in Atlantic salmon. Regardless, a natural question arises: how realistic is our experimental setting? Obviously, an IP injection does not reflect the actual route of infection, as it immediately exposes the fish to an unnaturally high infectious dose that bypasses the host’s primary defense mechanisms. However, aquaculture breeding for disease resistance has commonly relied on challenge tests using IP injection to infect the population under study, even in commercial programs (Gonen et al. 2015; Aslam et al. 2020; Robinson et al. 2023). In contrast to the acute disease seen in this challenge, naturally infected fish often become chronically infected with a slowly progressive disease. The immune defense tries to kill the bacteria and limit their spread by forming granulomas, but *R. salmoninarum* can escape by hiding inside white blood cells that phagocytose them. This trait in the bacteria is why antibiotic treatment and vaccine development against BKD are extremely challenging.

Several phenotypic proxies derived from the BKD challenge experiment were used in our study to investigate the underlying genetics, beginning with the estimation of relevant variance components. More specifically, the proxies used were the number of surviving days both as a continuous and a right–censored trait and the overall and N_50_ survival as binary ones. Overall, moderate heritability estimates were obtained (0.16-0.56), suggesting there is potential for applying selective breeding practices. This range of heritability values is consistent with values previously documented for various diseases in salmonids (Yáñez et al. 2014; Lhorente et al. 2019). It should be noted, though, that especially in the case of overall survival, the heritability estimate (0.26) was accompanied by a substantial standard error (0.15). Furthermore, the Gibbs sampler exhibited poor mixing and high autocorrelation, indicating that the posterior was sampled inefficiently and the resulting estimate is likely unreliable. This inefficiency stems from the high degree of skewness in the data, a condition under which the Gibbs sampler often fails to explore the posterior distribution effectively (Roberts and Sahu 1997).

In contrast to the moderate heritability estimates that were observed for surviving days and overall or N_50_ survival, a low heritability (0.054) was found for Ct values from RT-qPCR. Nevertheless, the latter estimate was based on only 800 records compared to 1982 for the other traits. In addition, it could well be that all those traits capture aspects of both disease resistance and resilience, which may differ in magnitude, thereby explaining the large differences between the heritability estimates. The relationship between survival and bacterial load, quantified through Ct values from RT-qPCR, yielded a weak negative Pearson correlation of −0.15. While lower Ct values indicate higher bacterial loads, the weak nature of this correlation suggests that the genetic mechanisms governing survival time may be partially distinct from those governing pathogen proliferation. This distinction is critical for breeding programs; selecting solely for "survival" may not necessarily result in "resistance" (the ability to limit bacterial load), and vice versa. In that aspect, breeding for disease resilience might be a more effective strategy (Knap and Doeschl-Wilson 2020).

The GWAS (Figure 5) provided the first insights into the genetic architecture of BKD resistance in Arctic charr, suggesting that BKD resistance is polygenic. It is worth noting that, with few exceptions, most aquaculture breeding studies reported polygenic inheritance in disease resistance (Robinson et al. 2023; Yáñez et al. 2023), favouring the application of genomic prediction over marker-assisted selection. The ability to accurately predict resistant individuals (AUC = 0.72) using the medium-density array (Figure 6) demonstrates that genomic selection is a viable tool for improving BKD resistance in Arctic charr. Incorporating these genomic breeding values into the national program, which currently relies on pedigree-derived breeding values, will enable the selection of resistant candidates lacking phenotypic data, thereby significantly increasing selection accuracy.

In light of the above, it is important to stress that before adjusting the overall breeding goals, it is pivotal to ensure that no unfavorable associations exist among the selected traits. In our particular case, the focus would be on assuring that selecting for increased BKD resistance would not negatively affect selection for increased growth. Unfortunately, since only a subset of the parental generation was genotyped, only a preliminary analysis was performed (involving 36 broodfish). The fact that only a slight and positive trend was observed amongst the breeding values of those two traits suggests that selecting for BKD resistance should not negatively affect selection for increased growth. However, to form a more complete picture, additional genotyping would be required.

## Ethics approval and consent to participate

All experiments on animals were carried out in strict accordance with the European guidelines and recommendations on animal experimentation and welfare. The disease challenge study was approved by the ethics committee on animal experimentation of the Norwegian Food Safety Authority, under the National Assignments Department, under No. 31329.

## Acknowledgments

The authors acknowledge support from Jordbruksverket in Sweden under the GenoAvel project (grant agreement 2023-646). GenoAvel was financed through the Maritime, Fisheries and Aquaculture Programme (EMFAF) 2021 - 2027, which is funded by both the EU and Sweden. The authors would also like to thank Benchmark Genetics for their collaboration during the SNP array development and genotyping.

## Authors’ contributions

CP conceived the study and conducted the genetic analysis. HH, ØE, and CA provided the bacterial strain and assisted with the challenge study. All authors contributed to drafting the manuscript. All authors read and approved the final manuscript.

